# The Wright-Fisher Site Frequency Spectrum as a Perturbation of the Coalescent’s

**DOI:** 10.1101/332817

**Authors:** Andrew Melfi, Divakar Viswanath

## Abstract

The first terms of the Wright-Fisher (WF) site frequency spectrum that follow the coalescent approximation are determined precisely, with a view to understanding the accuracy of the coalescent approximation for large samples. The perturbing terms show that the probability of a single mutant in the sample (singleton probability) is elevated in WF but the rest of the frequency spectrum is lowered. A part of the perturbation can be attributed to a mismatch in rates of merger between WF and the coalescent. The rest of it can be attributed to the difference in the way WF and the coalescent partition children between parents. In particular, the number of children of a parent is approximately Poisson under WF and approximately geometric under the coalescent. Whereas the mismatch in rates raises the probability of singletons under WF, its offspring distribution being approximately Poisson lowers it. The two effects are of opposite sense everywhere except at the tail of the frequency spectrum. The WF frequency spectrum begins to depart from that of the coalescent only for sample sizes that are comparable to the population size. These conclusions are confirmed by a separate analysis that assumes the sample size *n* to be equal to the population size *N*. Partly thanks to the canceling effects, the total variation distance of WF minus coalescent is 0.12*/* log *N* for a population sized sample with *n* = *N*, which is only 1% for *N* = 2 *×* 10^4^.

## Introduction

An attractive aspect of genealogical analysis is that it begins with current samples whose sequence data are directly measured. Wright-Fisher (WF) and the coalescent are two theoretical models used to make deductions about the genealogies of the current samples [Durrett, 2008].

The coalescent was derived and justified by Kingman [1982a] as an approximation of the WF model. Kingman’s analysis and extensions by other authors [Möhle, 2000, Möhle and Sagitov, 2001] assume the current sample size *n* to be fixed as the haploid population size *N* becomes large.

In view of the rapid increase in sample sizes in human genetics (see [Karczewski et al., 2016], for example), it is worth asking how close the WF and coalescent models are for large samples. A key property of the coalescent is that the genealogy is constructed entirely using binary mergers. In earlier work [Melfi and Viswanath, 2018], we have shown that for sample sizes *n* = *o* (*N* ^1*/*3^), WF genealogies involve only binary mergers with probability tending to 1. A more precise result derived here states that if the sample size is given by *n* = *αN* ^1*/*3^, the probability that the WF genealogy involves only binary mergers is approximately exp *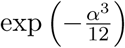*. To understand the onset of the deviation of WF from the coalescent, Fu [2006] as well as Bhaskar et al. [2014] looked at triple mergers, where three individuals merge into a common parent over a single WF generation. Among other results, we show that if the sample size is *n* = *αN* ^1*/*2^, the expected number of triple mergers in the WF genealogy is *α*^2^*/*6 + (exp(–*α*^2^*/*2) 1 */*3. This last result is in agreement with the *N* ^1*/*2^ scaling deduced in [Melfi and Viswanath, 2018].

With regard to sequence data, such results are perhaps too exacting. Detailed agreement in the genealogy is essential to reproduce the Kingman partition distribution [Kingman, 1982b] at each step of the genealogy. However, summary statistics such as the site frequency spectrum are not so refined. The site frequency spectrum of a sample of size *n*, which may be directly obtained from sequence data, consists of the probability that *j* of the samples are mutants and *n-j* are ancestral, *j* = 1, *…, n-*1, at a base pair. The site is assumed to be polymorphic with a single mutation at some individual that is an ancestor of some but not all samples.

The site frequency spectrum has been widely used for making demographic inferences (see [Excoffier et al., 2013, Fu et al., 2013, Griffiths and Tavaré, 1998, Gutenkunst et al., 2009, Kamm et al., 2017, Keinan and Clark, 2012, Lukic and Hey, 2012, Wakeley and Hey, 1997], for example) and is therefore a good basis to understand the difference between WF and the coalescent with regard to sequence data. If the genealogy is given by the coalescent and if *µ* is the mutation rate per site per generation, the probability that *j* out of *n* samples are mutants is

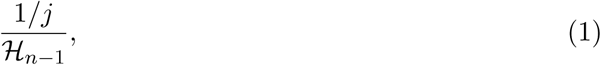

where *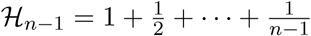* is the harmonic number, assuming *µ* to be so small that *µN* is negligible and assuming the sample to be polymorphic at the site. We will derive the first perturbing terms that follows (1) under the assumption of WF genealogy.

The elegant formula (1) for the probability of *j* mutants has a long and complicated history. Fisher [1922] stated that the correlation between heights of fathers and sons was 0.5 and attempted to obtain a Mendelian explanation of that correlation. He was thus led to a consideration of “gene ratios,” which is equivalent to counting the number of mutants. He derived the numerator of (1) some years later [Fisher, 1931]. Wright [1931, p. 120] had contacted Fisher earlier, noting (among other discrepancies) that he obtained 2*N* log(1.8*N*) for the size of the genealogy, whereas Fisher [1922] had obtained 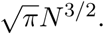^1^ There can be little doubt that Fisher was aided by [Wright, 1931] in coming up with the arguments that led him to the numerator of (1) as well as another result we will review shortly.

The size of the genealogy under WF is equal to the number of ancestors (with the current sample included) with 1, *…, n-*1 (but not *n*) descendants in the sample; in other words, the number of ancestors who would make the sample polymorphic if hit with a mutation. The *b*-branch length of the genealogy is the number of ancestors with exactly *b* descendants for *b* = 1, *…, n -*1. The size of the genealogy and the *b*-branch length are defined analogously for the coalescent, with the difference that the number of generations a lineage survives is no longer an integer. Ancestors of the current sample will be referred to as ancestral samples. Ancestors of the current sample in the same generation will be referred to as an ancestral sample. An ancestral sample induces a partition of the current sample, and for the coalescent, the partition follows the Kingman partition distribution [Durrett, 2008, Kingman, 1982b, Griffiths and Tavaré, 1998, Melfi and Viswanath, 2018].

Kimura [1955] (also see [Kimura, 1964, p. 222]) solved the diffusion equation for gene frequencies. From that point, (1) can be derived, although an argument connecting mutant frequencies in the population to that in the sample (such as the argument in [Durrett, 2008, p. 51]) would be needed. The first such argument was given by Ewens [1972], who also introduced the “frequency spectrum” terminology. A coalescent derivation of (1) was given by Fu [1995], as a consequence of the expectations and variances of *b*-branch lengths of the coalescent genealogy and was preceded by the treatment of special cases [Fu and Li, 1993, Tajima, 1989]. A mathematically complete treatment, allowing for varying population sizes, is due to Griffiths and Tavaré [1998].

If the genealogy is given by WF, we show that the probability of *j* mutants out of *n* is given by

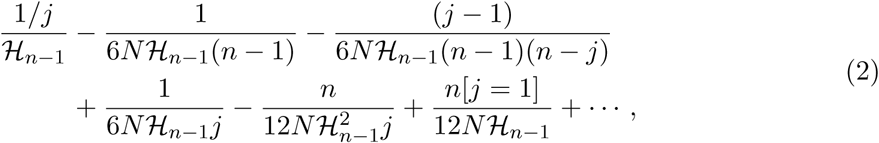

where [*j* = 1] is 1 if the assertion *j* = 1 is true and 0 otherwise and where *j* = 1, *…, n-*1. The result (2) is perturbative in that it gives the *N* ^*-*1^ terms but not the *N* ^*-*2^ terms.

Both (1) and (2) assume the population size *N* to be constant. Assuming the population size to be constant allows us to carry out an analysis and obtain information that would not be available from an algorithmic computation. The coalescent probability (1) does not depend upon *N*, although the probability of *j* mutants in the sample can be used to make demographic inferences. The WF probability (2) shows the dependence on *N*. Thus, limiting ourselves to constant *N* is not without advantages.

It is well-known that the first neglected term in a power series often gives a good idea of the error. In the same way, the 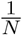 terms in (2) give an idea of the various phenomena at work in making the WF frequency spectrum differ from that of the coalescent. As noted by earlier authors [Bhaskar et al., 2014, Fu, 2006, Wakeley and Takahashi, 2003], the WF frequency spectrum elevates the probability of singletons (*j* = 1 mutants) and lowers the probability of *j* mutants for each *j >* 1. Such a movement in mutant probabilities may be verified explicitly from the last two terms of (2), which are the only terms that increase with *n*. The last two terms increase approximately linearly with sample size. For population sized samples, (2) yields an estimate of 1*/*12 *ℋ*_*N-*1_ (or 1*/*12 log *N*) for the amount by which singleton probability is raised under WF. Even with later terms in the perturbative series not taken into account, that estimate is off only by a factor of 3*/*2.

We give a separate analysis of population sized samples with *n* = *N*, with the work of Wakeley and Takahashi [2003] being our starting point. Fisher [1931] gave an ingenious derivation of *b*-branch lengths of WF genealogies with *n* = *N*, although some of his arguments are not entirely clear.^2^Wakeley and Takahashi [2003] gave a different and more transparent argument for the *b* = 1 case, which we extend to *b >* 1.

If *p*_*j*_and *q*_*j*_are two probability distributions over *j* = 1, *…, n–*1, the total variation (TV) distance between them is 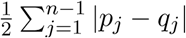 The total variation distance is the maximum difference in probabilities of any possible event under the two distributions [Brémaud, 1999, p. 126] and is therefore a quite robust way to compare probability distributions. The total variation distance between the frequency spectrums under WF and the coalescent for a population sized sample with *n* = *N* is approximately

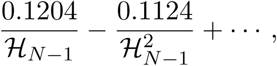

with a slight change in the approximation for *N >* 6.8 × 10^5^. For *N* = 2 × 10^4^, the baseline assumption in human genetics [Durrett, 2008], the total variation distance is only around 1%.

Fu [2006] has connected the greater speed of mergers under the coalescent to the elevation of singleton probability under WF. As we will explain, the coalescent is indeed faster for *n N* ^1*/*2^ but for *n≈ N*, the picture is not so clear. We refer to the same phenomenon as a mismatch in rates of merger to cover both cases. Another difference between the models is in the way children are partitioned between the parents [Melfi and Viswanath, 2018]. In particular, the offspring distribution is approximately Poisson for WF but approximately geometric for the coalescent.

To disentangle the two effects, we define an intermediate model called the discrete coalescent. In the discrete coalescent, the number of parents of a sample of size *n* has exactly the same distribution as in WF. However, once the number of parents is determined, the children are split between the parents according to Kingman’s partition distribution [Kingman, 1982b]. The intermediate model shows that the effect of the mismatch in rates is twice as great as the effect of the difference in the way children are split between parents. The two effects are of opposite sense and combine to cause a reduction in overall error.

## Poisson approximations to Wright-Fisher genealogies

In a backward WF step, each haploid individual chooses one out of *N* parents with equal probability and independently of all other individuals in its generation. The WF genealogy of a sample is built up using backward WF steps. The coalescent [Kingman, 1982a] may be thought of as a rate varying Poisson approximation of WF genealogies.

Other Poisson approximations may be used to capture more detailed information about WF genealogies. The clumping heuristic is a general method for deriving Poisson approximations [Aldous, 1989]. Applications of the heuristic require greater sophistication when the “clumps” are disconnected. In the case of WF, the clumps have a relatively simple form and the heuristic is not difficult to apply.

The following basic fact is all that we will need. Suppose the probability of occurrence of an event (such as a thunderstorm) in the interval (*u, u* + *du*) is *λ*(*u*) *du*. Then the total number of occurrences of the event in the domain [*a, b*] has Poisson distributed with rate 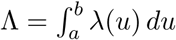 In particular, the probability of *k* occurrences is 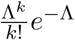 If an event is rare in every neighborhood, the total number of occurrences is approximately Poisson with the rate obtained by summing over the domain.

Let *n* be the number of samples and *N* the size of the parental generation. If *δ* is the number of samples lost due to mergers in a single backward WF step, the number of parental samples is *n - δ* and we have

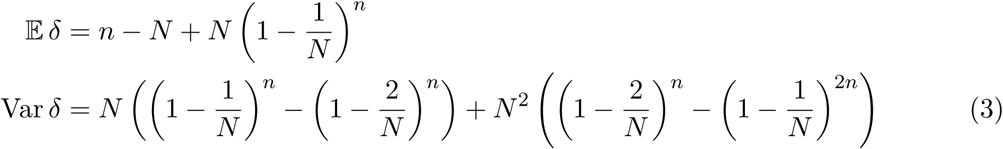

[Watterson, 1975]. When *n* is fixed, 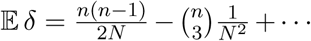 and Var *δ -*𝔼 *δ* = *-*2*n*^3^*/*3*N* ^2^ +*…*, suggesting a Poisson approximation for small *n* which turns out to be the Kingman coalescent. More generally, if *n* = *N* ^a^, where a *∈* [0,1), we have 𝔼 *δ -*Var *δ* = 𝒪 (*N* ^3a*-*2^) = *o* (*N* ^a^), suggesting a Poisson approximation to *δ* for a *<* 1.

For *i* = 1, *…, k*, the *i*th sample has the same parent as the *k* + 1st sample with probability 1*/N*, which is a rare event for *N≫* 1. The accumulated rate is *k/N*. Thus, the probability the *k* +1st sample has the same parent as one of the prior samples is approximately 1–exp(–*k/N*).

Suppose the number of samples is *n*. The probability that the *i* + 1st sample merges with one of the prior samples is 1 *-* exp(*-i/N*) approximately, which is a rare event for *i≪N*. For *n ≪ N*, the cumulative rate 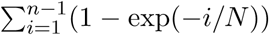 is the left hand Riemann sum of the integral

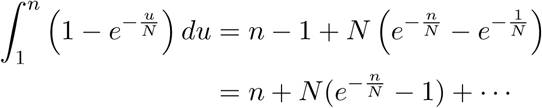

We may correct for an error that occurs in replacing the sum by an integral by taking *λ*_*δ*_(*n*) =*n* + *N* (exp(*-n/N*)*-*1)*-n/*2*N*. (The sum is now approximated to order *N* ^*-*2^.)

For *n≪N*, *δ* approximately follows a Poisson distribution of rate *λ*_*δ*_(*n*). Thus, ℙ (*δ* =*k*) ≈ exp(*λ*_*δ*_(*n*)) × *λ*_*δ*_(*n*)^*k*^*/k*!. In fact, 𝔼*δ* = *λ*_*δ*_(*n*) + *∈*, where *∈* = *n*^2^*/*2*N* ^2^ + is of the same order as the error in the Poisson approximation

Let Λ_2^2^_ (*n*) be the accumulated rate of occurrence of a double binary merger in some generation of the WF genealogy of a sample of size *n*. If the sample size is small enough, mergers in any generation are likely to be single binary mergers as in the Kingman coalescent. As the sample size increases, multiple binary mergers may appear with some likelihood and then triple mergers and so on [Melfi and Viswanath, 2018]. Using the Poisson heuristic illustrated above, it is shown in the Appendix (which has the same organization as the main text to facilitate looking up references to it) that

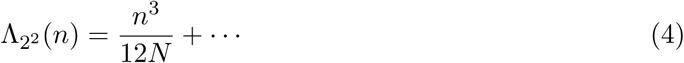

If *n* = *αN* ^1*/*3^, Λ_2^2^_ (*n*) = *α*^3^*/*12 implying the probability of coalescence with only binary mergers to be 1*-*exp *-α*^3^*/*12 (see Figure 1) and the probability of exactly *k* binary mergers in the genealogy to be exp(*-β*)*β*^*k*^*/k*!, where *β* = *α*^3^*/*12.

**Fig. 1:**
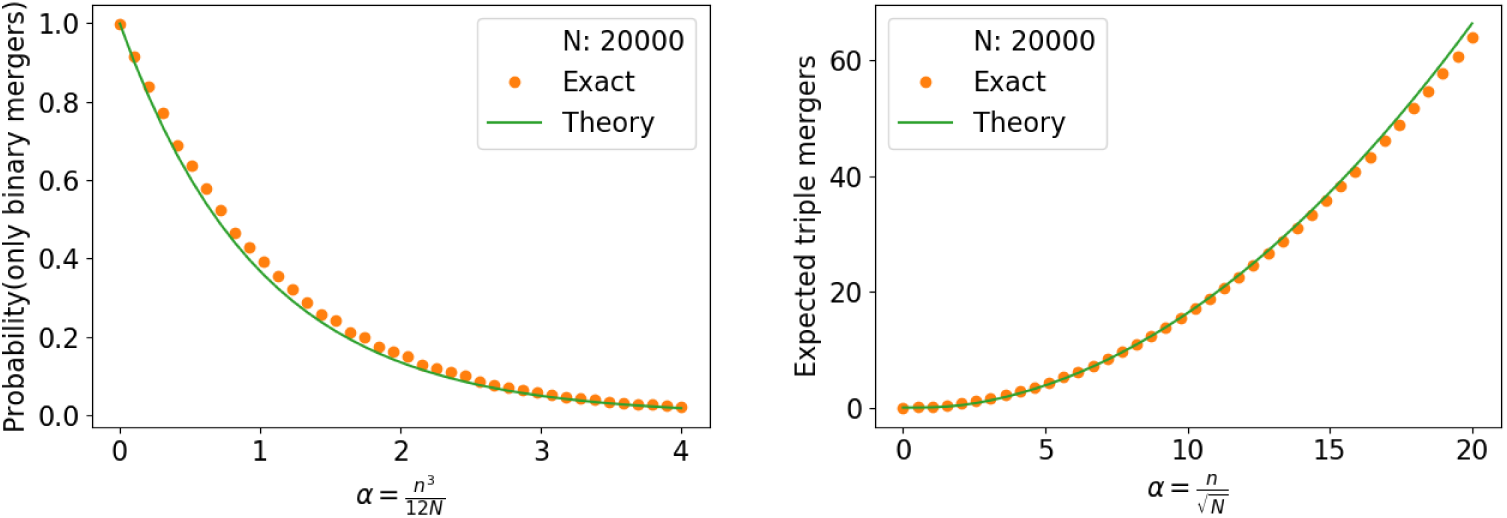
Plots verifying the approximations implied by (4) and (5). The exact numbers are from computer programs described in [Bhaskar et al., 2014, Melfi and Viswanath, 2018].

Let Λ_2_*p* (*n*) be the accumulated rate of occurrence of a *p*-fold binary merger in some generation of the WF genealogy of a sample of size *N*. Then (see Appendix)

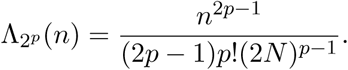

Thus, the correct scaling for the onset of *p*-fold binary mergers is 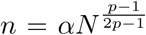 The scaling was given in earlier work [Melfi and Viswanath, 2018] but not the Poisson approximation.

For triple mergers, if 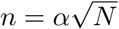, we have (see Appendix)

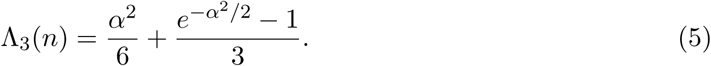

We may then use the Poisson distribution and approximate the expected number of triple mergers in the genealogy as Λ_3_(*n*) (see Figure 1) or calculate the probability of *k* triple mergers in the WF genealogy of the sample. The *N* ^1*/*2^ scaling of triple mergers was determined in earlier work [Melfi and Viswanath, 2018].

## Perturbative analysis of the WF site frequency spectrum

The manner in which coalescent and WF genealogies differ may be inferred from (4), (5), and other similar results. Such differences in genealogy are a part of modeling and are not directly observable from sequence data. The question becomes to what extent the genealogical differences show up in sequence data.

In this section, we will outline the the main ideas in obtaining the WF frequency spectrum. The leading term of course is the coalescent answer, which is 1*/j ℋ*_*n–1*_. We will calculate the following *N* ^*-*1^ terms. In this section, we give an overview of the analysis with a focus on the *j* = 1 singleton case. The Appendix treats the general *j* case and supplies the details.

The first perturbing terms, which we will calculate, suggests that the correct scaling for the divergence of WF frequency spectrum from that of the coalescent is *n* = *αN*. Although not a proof, the suggestion is almost surely correct and we verify it from another angle later. The scaling for the onset of simultaneous binary mergers and triple mergers is *N* ^1*/*3^ and *N* ^1*/*2^ (from (4) and (5)). The fact that the divergence in the frequency spectrum sets in for much larger samples means that the frequency spectrum is not very sensitive to multiple mergers in the genealogy.

If the WF genealogy of a sample of size *n* progresses through sample sizes as in

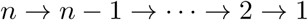

without skipping any sample size in-between *n* and 1, we denote that no-skip event by 𝒮 _0_. If the WF genealogy skips from a sample size of *m* + 2 to *m*, omitting *m* + 1, we denote such a skip-to-*m* event by *𝒮* _*m*_ for *m* = *n -* 2, …, 1. The sample size of *m* + 1 is the only omission in 𝒮*m.*

Other patterns of skipping are possible. However, the probability of such events is 𝒪 (*N* ^*-*2^). For an 𝒪 (*N* ^*-*1^) calculation, we only need to consider 𝒮 _0_ and 𝒮 _*m*_.

The WF frequency spectrum is calculated under the assumption of exactly one mutation in the genealogy of the sample. Therefore, we define the event 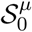 to be *𝒮* _0_ and exactly one mutation in the genealogy of the sample. The event 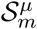 is defined analogously.

## Coalescent and WF propagators

The general approach to derive the WF frequency spectrum is to first determine the probability that a mutation occurs in the genealogy when the sample size is *m* for *m* = *n*, …, 2. That would mean that 1 out of *m* ancestral samples is a mutant at some point in the genealogy. That probability is then propagated to the current sample size of *n*.

If the genealogy from the current sample size of *n* to the ancestral sample size of *m* involves only binary mergers, the probability of a single mutant in the sample of *n* given a single mutant in the ancestral sample is

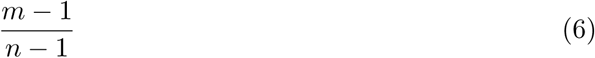

(see Appendix). The Appendix derives this formula as a special case of a more general formula for propagating probabilities under the coalescent.

To find the first perturbing term in the WF frequency spectrum, there is only one more case to consider. Suppose that a sample of size *m* + 2 changes into a parental sample of size *m* under a single backward WF step. Given that the parental sample of *m* has only a single mutant, the probability that the sample of size *m* + 2 has only a single mutant is

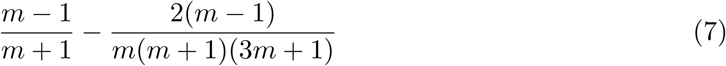

(see Appendix). Thus, skipping a step under WF reduces the factor that propagates the probability of a single mutant.

### Probability of mutation at *m*

What is the probability of a mutation event at *m* assuming that the sample size *m* is visited? Consider the following picture:

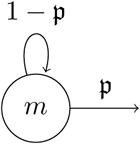

The picture is showing that an ancestral sample size of *m* remains *m* under a backward WF step with probability 1 𝒫 and exits to a lower sample size with probability 𝒫. Neglecting *µ*^2^ terms, the probability that a sample of size *m* will be hit with a mutation is

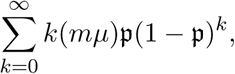

where *k* is the number of returns from *m* to *m*.

Thus, with *µ*^2^ terms neglected, the probability of being hit with a mutation at *m* is equal to *mµ/* 𝒫. By plugging in a suitable value for 𝒫 (see Appendix), we get the probability of being hit with a mutation at *m* to be

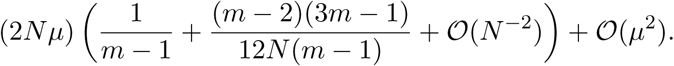

Neglecting *µ*^2^ and *N* ^*-*2^ terms, we denote the probability of being hit with a mutation at *m* by

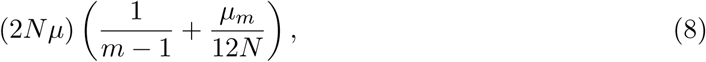

where *µ*_*m*_ = (*m-*2)(3*m-*1)*/*(*m-*1) + 𝒪 (*N* ^*-*1^).

In fact, because we are neglecting *µ*^2^ terms, (8) gives the probability that there is a single mutation in the entire genealogy with that mutation occuring when the ancestral sample size is *m*.

### Probability that *m* + 2 **skips to** *m*

Suppose the ancestral sample size is *m* + 2. What is the probability that the ancestral sample size skips over *m* + 1 and goes directly to *m* under WF? The ancestral sample size could skip over both *m* + 1 and *m*, but because we are neglecting *N* ^*-*2^ terms, those possibilities may be ignored.

The probability of skipping to *m* is (see Appendix)

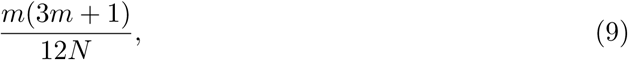

with *N* ^*-*2^ terms neglected.

### The event *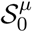*

From (9), it follows that that the probability of *𝒮*_0_, which visits each ancestral sample size in {1, *… n*} is 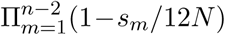 Using (8), the probability of a single mutation in the genealogy is (2*N µ*)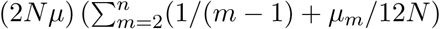, with *µ*^2^ terms ignored and with *N* ^*-*2^ terms ignored in the coefficient of 2*Nµ*. The summation over *m* = 2, *…, n* sums over the probability of the single mutation occurring when the ancestral sample size is one of 2, *…, n*.

Thus, the probability of 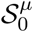 is 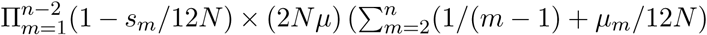. Simplifying and omitting *N* ^*-*2^ terms in the coefficient of 2*Nµ*, we get 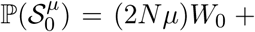 with

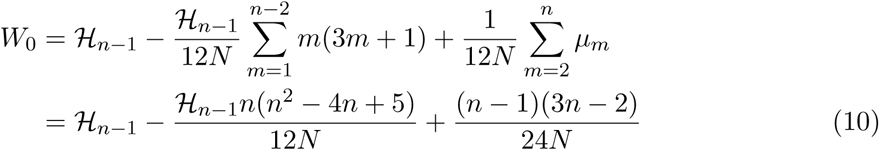

and with *N* ^*-*2^ terms neglected in *W*_0_. The last step in (10) is gotten after a routine simplification. At this point, we can think of 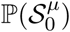 as proportional to the weight *W*_0_.

Let ℳ_*m*_ be the event that a mutation occurs in the genealogy of the sample of size *n* when the ancestral sample size is *m*. From (8), we know that the probability that a mutation occurs at *m* but nowhere else in the genealogy is proportional to 1*/*(*m -* 1) + *µ*_*m*_*/*12*N*. Therefore,

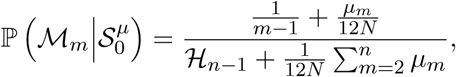

where the denominator is obtained by summing over *m* = 2, *…, n*. The right hand side above can be simplified to obtain

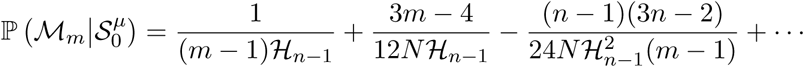

with *N* ^*-*2^ terms ignored and in the limit *µ* 0.

The number of mutants in the current sample of size *n* is always denoted by *j*. The next step is to calculate 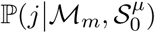 For *j* = 1, we can use (6) to propagate a single mutant from an ancestral sample of size *m* to the current sample of size *n* and get

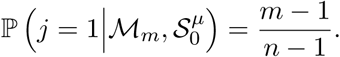

The general *j* case is treated in the Appendix. Finally, we can calculate 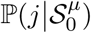 using

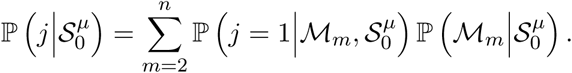

At that point the analysis of the 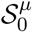 case is complete.

### The event *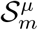*

From (9), ℙ (*S*_*m*_), which is the probability the genealogy skips from sample size *m* + 2 to *m*, is

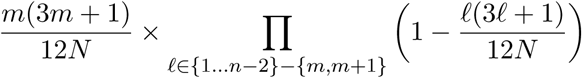

or simply *m*(3*m* + 1)*/*12*N* with *N* ^*-*2^ terms neglected.

Because ℙ (*S*_*m*_) leads with a *N* ^*-*1^ term, we may simplify (8) and take the probability that a mutation hits when the ancestral sample size is *l* to be (2*N µ*)*/*(*l -* 1). It follows that 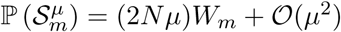

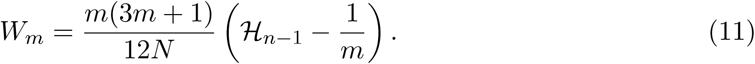

with *N* ^*-*2^ terms neglected in *W*_*m*_. At this point, we can take 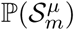 to be proportional to *W*_*m*_.

Let ℳ_2*…m*_ denote ℳ_2_⋃… ⋃*ℳ*_*m*_, in words, the event where a mutation occurs when the ancestral sample size is 2, *…, m*. The probability that a mutation strikes when the sample size is *l* is proportional to 1*/*(*l -* 1). Therefore,

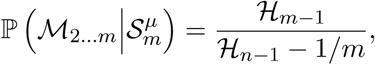

with all *N* ^*-*1^ and *µ* terms ignored. The probability of a single mutant in the ancestral sample of size *m* given *ℳ*_2*…m*_ is 1*/ℋ*_*m-*1_ (by (1)) with *N* ^*-*1^ terms ignored. That probability propagates to *m* + 2 with the factor (7) and from *m* + 2 to *n* with the factor 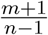 (by (6)). Therefore, with *N* ^*-*1^ terms ignored and in the limit *µ ⟶* 0, we have

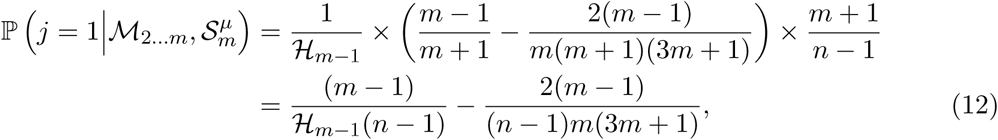

where the second term accounts for the WF correction for the non-binary merger that occurs in skipping to *m*. The calculation of 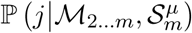 in the general *j* case is given in the Appendix. The analysis of 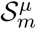 is complete once 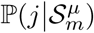 is obtained.

The WF frequency spectrum (2) is obtained by simplifying (see Appendix)

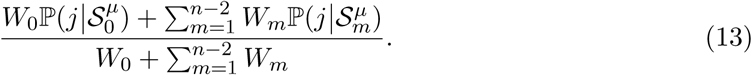

If we look at the sequence steps building up to this point, the difference in the way WF and the coalescent partition children between parents first comes up in the latter term of (7). That terms propagates to the latter term of (12). If we separate such terms in (13), we will have obtained the effect of differing partition distributions on the WF frequency spectrum.

The part of the *N* ^*-*1^ terms in the WF frequency spectrum (2) that is due to differences in partitioning between WF and the coalescent is given by (see Appendix)

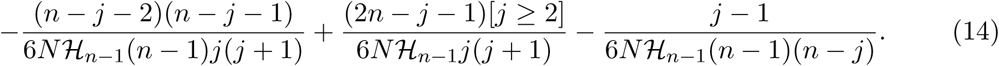

Evaluating with *j* = 1 and retaining only the dominant term, we get –*n/*12*Nℋ* _*n*_*-*_1_ to be the effect on singleton probability of the difference in partitioning distributions. Evaluating (2) with *j* = 1 and retaining only the dominant term, we obtain *n/*12*N ℋ*_*n* 1_ as the amount by which the WF singleton probability exceeds that of the coalescent. Therefore, the effect of the mismatch in rates of merger (as defined in the Introduction) must be *n/*6*N ℋ*_*n*_–_1_.

Figure 2 shows that the WF singleton probabilities are elevated and the rest of the frequency spectrum is depressed, as may be inferred from the last two terms of (2). The figure also illustrates the correction due to rates being twice as high as the correction due to differences in the way children are partitioned between parents.

**Fig. 2:**
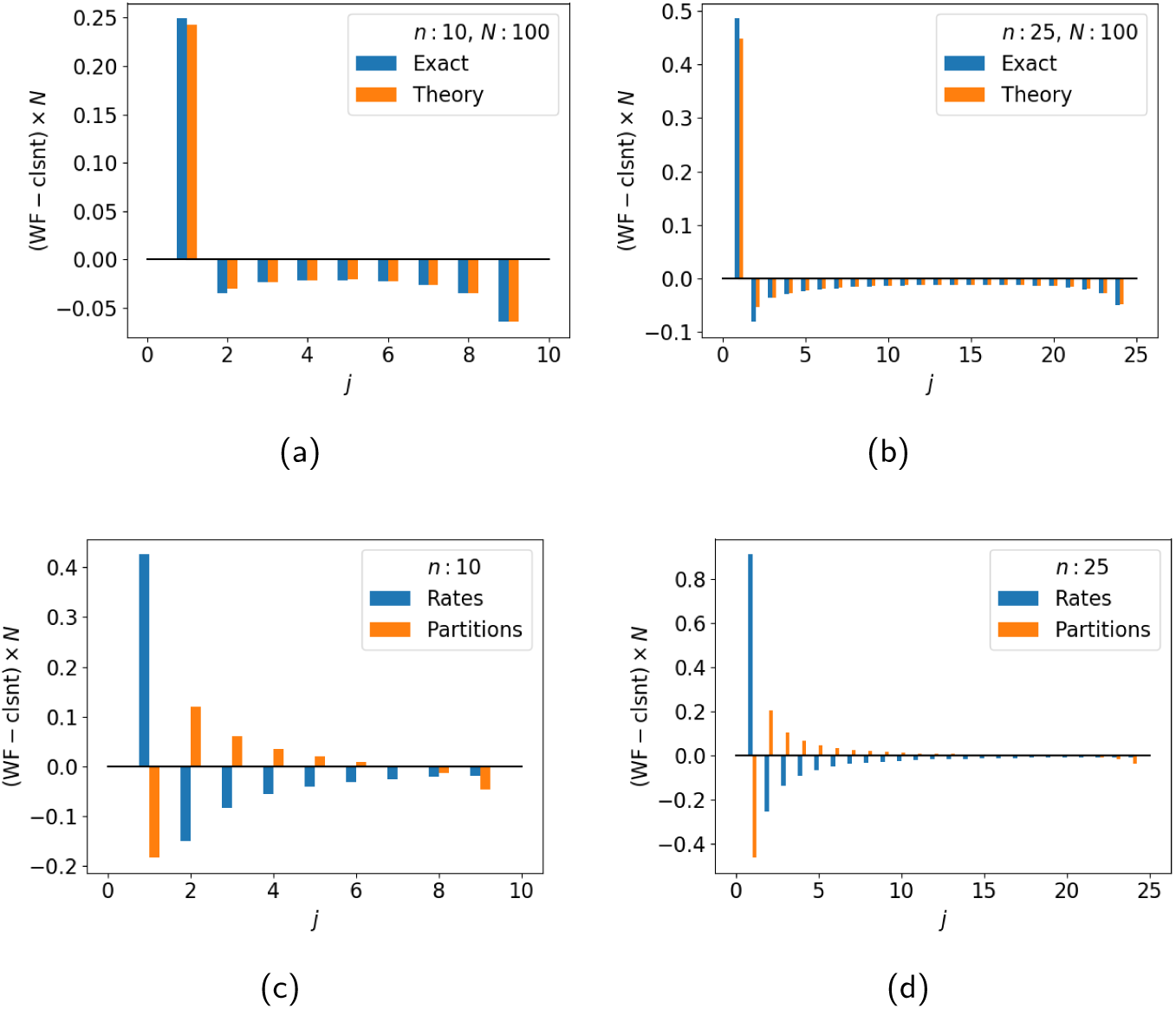
(a) and (b): WF minus coalescent computed using (2) minus (1) (theory) is compared with a computation using the program of Bhaskar et al. [2014] (exact). (c) and (d): (2) minus (1) minus (14) (rates) is compared with (14) (partitions).

Because *j* = 1 singleton probabilities are elevated under WF and other probabilities are lowered, we may obtain the total variation distance between WF and the coalescent by simply taking the difference in *j* = 1 probabilities. Thus, the perturbative estimate for the total variation distance between the WF frequency spectrum and that of the coalescent is *n/*12*Nℋ*_*n* 1_. This estimate is qualitatively correct even for *n* = *N*, and even quantitatively it is not unreasonable, being about 2*/*3rds of a better estimate we will presently derive. Figure 3 shows that the total variation distance increases with *n* and decreases with *N*.

**Fig. 3:**
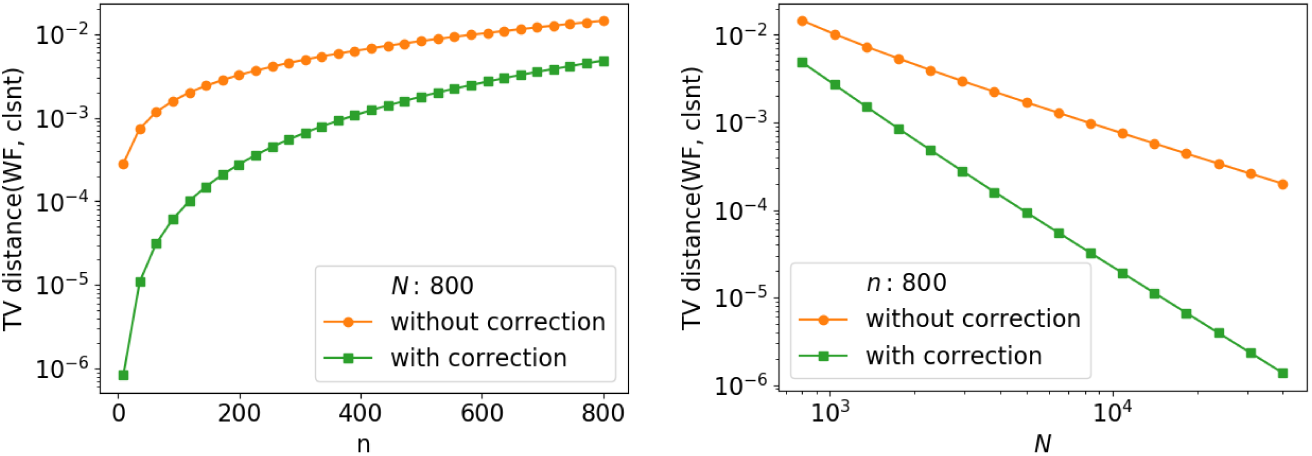
Total variation distance between WF frequency spectrum [Bhaskar et al., 2014] and that of the coalescent given by (1) (without correction) or with correction as given by (2).

## Population sized samples

Suppose the sample size is *n* = *αN*. For an individual among the parental population of *N*, the probability that any given sample is a child is 1*/N*, a rare event. The accumulated probability over the sample of size *αN* is *α*. Therefore, by the Poisson clumping heuristic, we may approximate the number of children of an individual in the parental generation by the Poisson distribution with rate *α*. The probability that an individual has *k* children among the *αN* samples is approximately exp(*-α*)*α*^*k*^*/k*!. The generating function 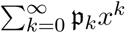 with 𝒫_*k*_ = exp(*-α*)*α*^*k*^*/k*! is exp(*α*(*x* 1)).

The only individuals in the parental generation that appear in the genealogy are ones who have at least one child among the samples. Therefore, it is natural to look at the Poisson distribution under the condition of having one child. Under that condition, the probability of having *k* children is 𝒫_*k*_*/*(1–exp(–*α*)) and the generating function is (exp(*αx*)–1)*/*(exp(*α*)–1).

Let *G*_1_(*αN*) = *g*_1_(*α*)*N* be the expected *b*-branch length with *b* = 1 of the WF genealogy of a sample of size *αN*. By (3), the expected number of parents is *N* (1–exp(–*α*)). Thus, we may write

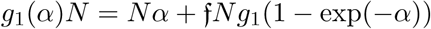

because the current samples *Nα* all contribute to the 1-branch length and with the understanding that f is the probability that a branch with a single descendant in the genealogy of the parental sample of size (1–exp(–*α*))*N* remains a branch with a single descendant in the genealogy of the current sample of size *αN*. That probability 𝒻 is the same as the probability of a parent having a single child, which is *α/*(exp(*α*) *-* 1). Therefore, we have

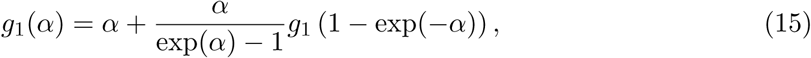

which is a result of Wakeley and Takahashi [2003] derived essentially using their arguments.

The generating function for the number of children of *a* parents is approximately

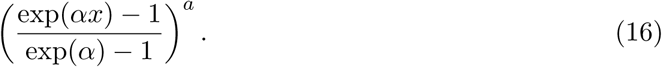

Therefore (see Appendix), the probability that *a* parents have *b* children is

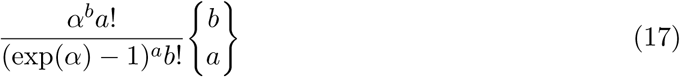

for *b* = *a, a* + 1, *…* Here 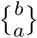 is a Stirling number of the second kind [Graham et al., 1994]. Using the same argument as above and taking the *b*-branch length with *αN* samples to be *G*_*b*_(*αN*) = *Ng*_*b*_(*α*), we get the recurrence

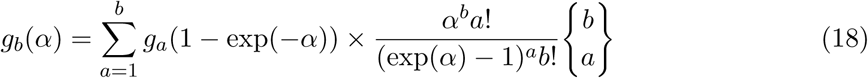

for *b* = 2, 3, *…*

By solving the recurrences for *g*_*b*_(*α*) and taking *α* = 1, we can obtain approximations to the WF frequency spectrum with *n* = *N* and compare it to (1), which is the coalescent frequency spectrum. However, we seek to separate the difference into a part due to the mismatch in rates of mergers and a part due to the difference in the way children are partitioned among parents.

We turn to the discrete coalescent, which is a model intermediate between the coalescent and WF. To obtain the manner in which *αN* children are split between *βN* parents under the discrete coalescent, which uses the Kingman partition distribution, we may fix an orange at the left most position and permute *βN-*1 identical oranges and *αN -βN* identical apples after it. The number of children of the *i*th parent can be taken to be the number of apples between the *i*th and *i* + 1st orange plus one (thus counting the *i*th orange) [Durrett, 2008, Griffiths and Tavaré, 1998, Melfi and Viswanath, 2018].

The probability that a parent has *k* children is approximately *γ*(1 − *γ*)^*k*^, with *γ* = *β/α* = (1 exp(–*α*))*/α* for a sample of size *αN*. The generating function of this geometric distribution is *γx/* (1 (1 *γ*)*x*). The generating function for the number of children of *a* parents is approximately (*γx/* (1 (1 *γ*)*x*))^*a*^. By extracting the coefficient of *x*^*b*^, we find the probability of *a* parents having *b* children under the discrete coalescent to be

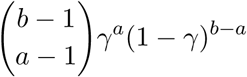

approximately.

If 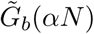 denotes the *b*-branch length of the discrete coalescent genealogy of *αN* samples, we may set 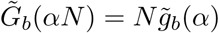 and obtain the recurrences

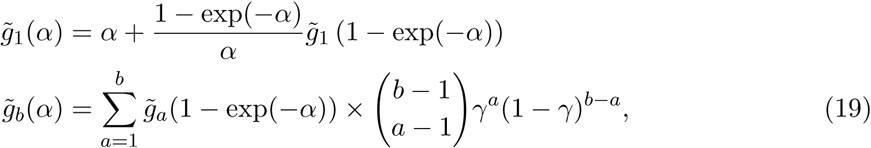

where *b* = 2, 3, *…*

In the Appendix, we show how to solve (15), (18), and (19) accurately using Chebyshev polynomials. For the coalescent, the expected *b*-branch length for a sample of size *n* = *N* is 2*N/b*. Therefore, we set *g*_*b*_(1) = (2*/b* + *ϵ*_*b*_)*N* and 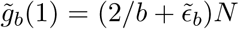 and report *∊*_*b*_ and 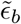 in Table 1.

**Table 1:** The expected *b*-branch length of the WF genealogy of *n* = *N* samples is (2*/b* + *∈*_*b*_)*N*. For the discrete coalescent, whose merger rates match WF but which partitions children between parents like the coalescent, it is 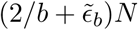.

| $b$ | $\epsilon_b$ | $\tilde{\epsilon}_b$ |
| --- | --- | --- |
| 1 | 0.240917257 | 0.418035261 |
| 2 | -0.046223840 | -0.100136471 |
| 3 | 0.005196946 | -0.032826669 |
| 4 | 0.001095702 | -0.017086273 |
| 5 | -0.000238278 | -0.011181848 |
| 6 | -0.000114882 | -0.008036411 |
| 7 | -0.000004091 | -0.006053860 |

The first column of the table agrees very well with Fisher [1922, p. 214]. To obtain the total size of the WF genealogy of *n* = *N* samples, we use

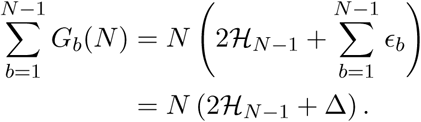

We estimate Δ to be 0.200645075 by summing *ϵ*_*b*_ over 1≤*b≤*20. The size of the discrete coalescent genealogy is the same as that of WF genealogy by definition. Our value for Δ agrees with Fisher’s except in the last decimal place.

The probability of *j* mutants in the WF spectrum of *n* = *N* samples is estimated to be

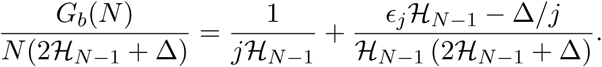

The estimated probability of *j* = 1 under WF exceeds 1*/jℋ*_*N–*1_ because *ϵ*_1_ *>* Δ. For *j* = 3, the term *ϵ*_*j*_*ℋ*_*N–*1_–Δ*/j* is negative as long as *N <* 6.8 × 10^5^but flips sign around *N* = 6.8 10^5^. For *j* = 4, *ϵ*_*j*_*ℋ*_*N–*1_–Δ*/j* turns positive only around *N* = 10^20^. Thus, with minor caveats, the WF frequency spectrum is elevated at *j* = 1 but depressed slightly for *j >* 1.

We can approximate the total variation distance between WF and coalescent frequency spectrums for *n* = *N* as

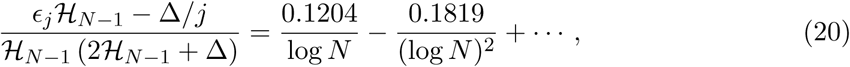

a result that is a direct consequence of [Fisher, 1922]. The first p lot o f F igure 4 s hows this estimate to be quite good. The figure also shows *g*_1_(*α*) is quite accurate for even *N* = 100 and small α, although *g*_4_(α) has visible errors for *N* = 100.

From Table 1, it is evident that the excess of the discrete coalescent’s *j* mutant probability over that of the coalescent

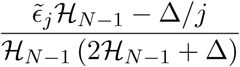

is positive for *j* = 1 and negative for *j >* 1. The discrete coalsecent and the coalescent differ only with respect to their rates of merger. Both of them follow the Kingman partition distribution. The effect due to difference in the rates of merger alone is twice as great because 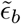 is nearly twice *ϵ*_1_.

When *n ≪N* ^1*/*2^, we can say that the coalescent is faster than WF [Fu, 2006] because *n*(*n -* 1)*/*2*N ≥* 𝔼*δ* (see Appendix). However, when *n≫N* ^1*/*2^, there can be several mergers in the same generation and rates of merger cannot be compared so directly. Although, the coalescent begins with a higher rate it adjusts its rate downwards with every binary merger.

During a single backward WF step a sample size of *n* =α*N* changes on an average to *m* = (1*-*exp(*-*α))*N*. On an average the coalescent takes 2*N* (1*/m-*1*/n*) generations to go from *n* samples to *m*. In fact, 2((1–exp(–α))-1–α^-1^) = 1 + –α*/*6+*… >* 1 (see Appendix), and the coalescent is in fact slower.

However, the 1-branch length of the coalescent in going from *n* samples to *m* is equal to 2*N* (*n -m*)*/*(*n-*1). It may be shown that 2*N* (*n-m*)*/*(*n-*1) *< N α*when *n* = *Nα* and *m* = *N* (1–exp(–*α*)) (see Appendix). Therefore, although the coalescent may take a little more than a generation to go from *n* samples to *m*, its 1-branch length is lower as a consequence of repeated binary mergers over slightly more than a generation.

**Fig. 4:**
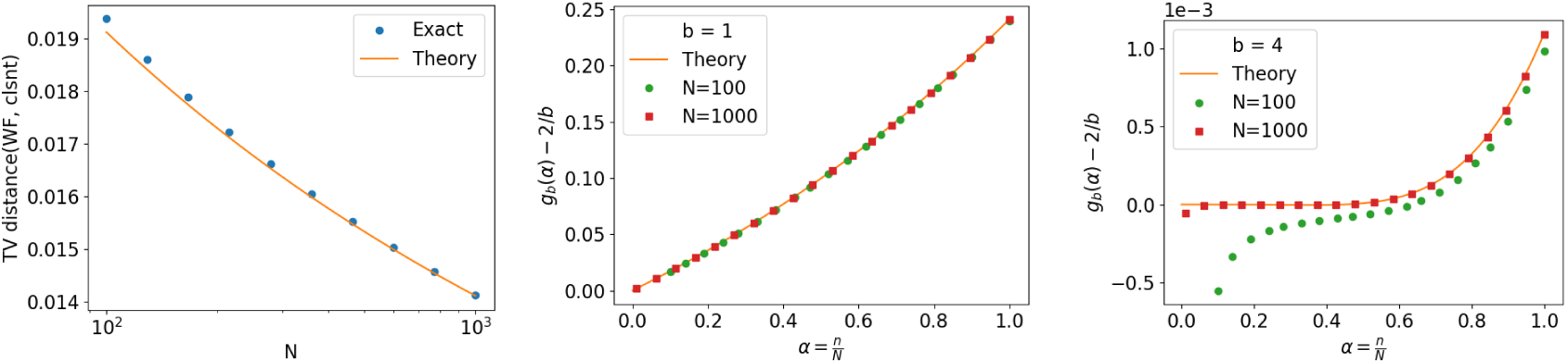
The first plot demonstrates the accuracy of (20). The next two plots examine the accuracy of *g*_*b*_(*α*). In all cases, the exact computations use the computer program of Bhaskar et al. [2014].

## Discussion

WF deviates from the assumptions of the coalescent for even small sample sizes. Simultaneous binary mergers appear in WF genealogies for sample sizes of only *αN* ^1*/*3^ with appreciable probability. Triple mergers appear for samples sizes of *αN* ^1*/*2^.

However, the effect of such deviations on the site frequency spectrum is minimal. Deviations in the site frequency spectrum set in only for sample sizes *αN*. Even for population sized samples the deviation is only around 1%. The effect is so small because the coalescent is self-correcting. The rate of mergers under the coalescent is faster, but the coalescent lowers the rate with every merger. The coalescent limits itself to binary mergers. As a result, the offspring distribution under the coalescent is geometric, whereas it is Poisson under WF. The geometric and Poisson distributions are not far enough apart to cause a major effect. In addition, the effect of differing offspring distributions partly cancels the effect of differing rates of merger.

Population substructure is perhaps the major reason to look for more sophisticated models than the coalescent [Durrett, 2008, Wakeley, 2009]. Skewed offspring distributions are another reason [Eldon and Wakeley, 2006, Matuszewski et al., 2018]. In the setting of skewed offspring distributions, it is known that the skew has to be comparable to the population size for deviations to show up [Eldon and Wakeley, 2006]. Thus, in that setting too, the coalescent is a robust model.

It is know that increasing skewness of offspring distribution raises the probability of singletons and lowers the probabilities of *j* mutants for *j >* 1 [Eldon and Wakeley, 2006]. The Poisson offspring distribution of WF has a lower variance that the geometric offspring distribution of the coalescent (see Appendix). Our finding that the effect of differing offspring distributions is to lower the singleton probability under WF is consistent with this point of view.

As far as the single site frequency spectrum is concerned, the coalescent is a robust and reliable model relative to WF. What if multiple sites are considered, possibly allowing for recombination between sites? Our conjecture is that the total variation distance between WF and the coalescent will still be of the order *C/* log *N* for even population sized samples. However, the constant *C* may increase with the number of sites. In that regard, we mention the availability of software to efficiently simulate WF genealogies under very general conditions [Palamara, 2016].

## Appendix

This appendix has the same organization as the main text. Corresponding sections of this appendix supply mathematical arguments omitted from the main text.

### Poisson approximations to Wright-Fisher genealogies

Mergers during a single backward WF generation occur at the *λ*_*δ*_(*n*). The number of samples lost due to merger is approximately Poisson with rate *λ*_*δ*_(*n*) = *n* + *N* (exp(*-n/N*) *-* 1) *-n/*2*N* for *n ≪N*.

Therefore, the probability of something other than a binary merger conditional on *δ ≥* 1 is

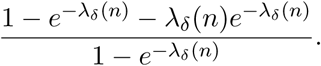

For *n≪N* ^1*/*2^, *λ*_*δ*_(*n*) = *n*^2^*/*2*N* is a good approximation. Because non-binary mergers at onset are double binary mergers [Melfi and Viswanath, 2018], we have the cumulative rate of double binary mergers (or non-binary mergers) to be

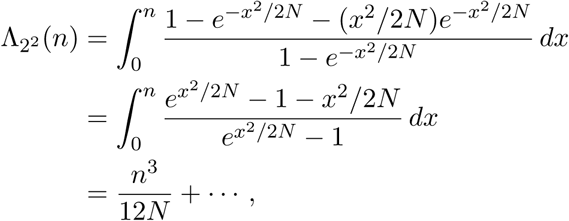

where the last step is from a power series expansion of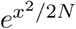.

The rate Λ_2^*p*^_ (*n*) for *p* simultaneous binary mergers is obtained similarly. In this case, a *p* simultaneous binary merger occurs during a single backward WF step conditional on *δ ≥*1 with probability

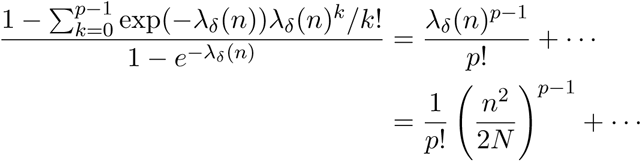

The accumulated rate over the entire genealogy is

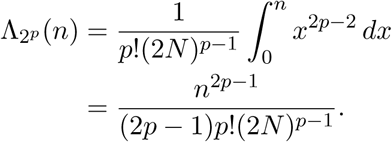

When multiple mergers or simultaneous binary mergers occur, the WF genealogy skips over sample sizes. We could correct for that, but the correction will not affect the leading term because we are still operating in the *n ≪N* ^1*/*2^ regime.

The reasoning for triple mergers is slightly different. We first need to obtain the rate of triple mergers during a single backward WF step. Consider the *m* + 1st sample. Each of the first *m* samples has the same parent as the *m* + 1st sample with probability 1*/N*, a rare event. Thus, the number of samples out of the first *m* that have the same parent as the *m* + 1st sample is Poisson with rate *m/N*. The probability that two of them have the same parent as *m* + 1, causing a triple merger, is

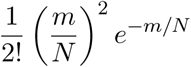

approximately. Therefore, the accumulated rate of triple mergers over a single generation is

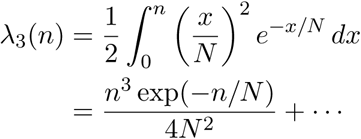

Triple mergers are a rare event for *n≪N* ^2*/*3^. However, when accumulating the rate of triple mergers over the entire genealogy, it is essential to account for the WF genealogy skipping some sample sizes.

The expected *δ* when the sample size is *n*, given that *δ ≥*1, is *λ*_*δ*_(*n*)*/* (1–exp(–*λ*_*δ*_(*n*)). Thus, for *m≤n*, we take the probability that *m* is reached to be (1–exp(–*λ*_*δ*_(*m*))) */λ*_*δ*_(*m*).

For the accumulated rate of triple mergers, we obtain

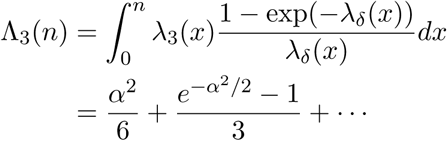

for *n* = *αN* ^1*/*2^ after using the approximation *λ*_*δ*_(*n*) = *n*^2^*/*2*N*.

### Perturbative analysis of the WF site frequency spectrum

The events *𝒮*_0_, *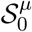, 𝒮*_*m*_, and 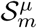 are assumed to be defined as before. We will derive (2), the *j* = 1 case of which was discussed in the main text. As before, *j* is always the number of mutants in the current sample of size *n*, which is assumed to be polymorphic with a single mutation in its genealogy.

#### Coalescent and WF propagators

Suppose an ancestral sample of size *m* has *i≥*1 mutants and (*m-i*) non-mutants. The genealogy from the current sample of size *n* to the ancestral sample of size *m* is assumed to involve only binary mergers, with no mutations in-between. As we will presently show, the probability that the current sample has *j ≥i* mutants is given by

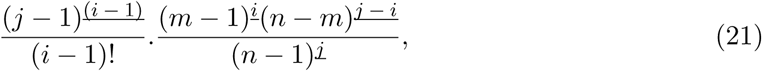

where *j*^*<u>i</u>*^ is the falling power *j*(*j-*1) *…* (*j-i* + 1) [Graham et al., 1994]. We adopt the convention that *j*^<u>0</u>^ is 1, even for *j* = 0. The coalescent propagator given earlier (6) is a special case (21) with *i* = *j* = 1.

The Kingman partition distribution may be used to obtain (21). However, we will give a more direct argument. Suppose an ancestral sample of size m has i mutants. Suppose that an ancestral sample of size m + 1 is related to it through a single binary merger. Then the probability that the sample of m + 1 has i + 1 mutants is i*/*m because each sample out of m is equally like to “split” and one of the i mutants will split with probability i*/*m. Similarly, the probability that the sample of m + 1 has i mutants is (m– i)*/*m (in this case, one of the m– i non-mutants has to split).

From here, we can write down the probability that a sample of *n* has *j* mutants when it is descended through binary splits from a sample of size *m* with *i* mutants to be

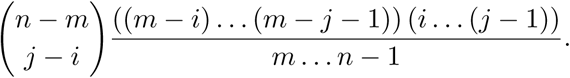

The argument for this expression is as follows. There are *n–m* splits from *n* to *m*. The binomial coefficient chooses *j–i* of those splits to be ones that increase the number of mutants. The denominator of the fraction in the expression steps from the sample size of *m* to the sample size of *n–*1 because those are the sample sizes that split. The numerator has the factor (*m-i*) *…* (*m–j -*1) to account for splits of non-mutants. The other factor *i …* (*j-*1) accounts for splits of mutants. The above expression is simplified to obtain (21).

When *i* = 1, (21) reduces to

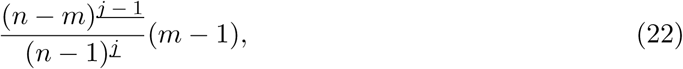

a useful special case.

Suppose next that a sample of size *n* is descended from a sample of size *m* with *i* mutants through a single backward WF step. It is assumed that there are no mutations during this descent. The probability of *j* mutants in the current sample is then given by

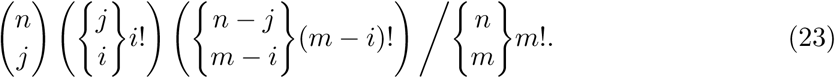

That is because in the current sample, we can choose *j* individuals to be mutants in 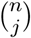 ways. That being done, the *j* mutants in the current sample can be assigned to *i* mutants in the parental sample, with each parent receiving at least one child, in 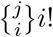 ways: the *j* samples can be partitioned into *i* in 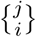 ways and then can be permuted in *i*! ways. The last bracketed factor in the numerator is the number of ways to assign (*n–j*) not mutants to (*m–i*) non-mutants in the parental sample. The denominator is the number of ways to assign *n* children to parents, with each parent receiving at least one child.

The Stirling numbers (of the second kind) 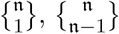, and 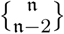 are given by 1, n(n *-* 1)*/*2, and n(n *-* 1)(n *-* 2)(3n *-* 5)*/*24, respectively [Graham et al., 1994]. Using (23) along with those formulas, we obtain the probabilities that a sample of size *m* + 2 has *i, i* + 1, *i* + 2 mutants when it is descended from an ancestral sample of size *m* in a single WF generation to be

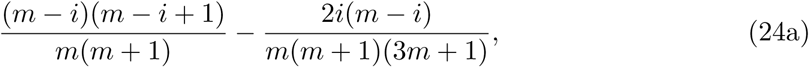

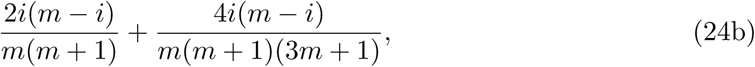

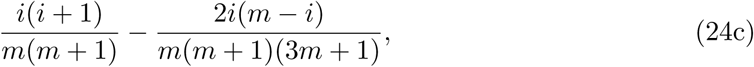

respectively. We may obtain (7) from (24a) by setting *i* = 1.

#### Probability of mutation at *m*

In the corresponding part of the main text, 𝒫 is the probability that a backward WF step applied to a sample of size *m* will reduce the sample size. We may take

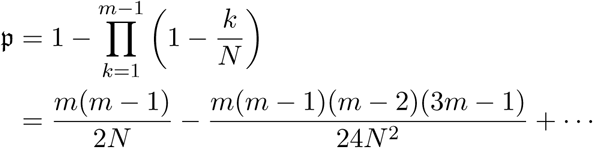

by ignoring terms after *N* ^*-*2^. In the main text, we showed that the probability of being hit with a mutation at *m* to be *µm/*𝒫. We may derive (8) by substituting for 𝒫 in *µm/*𝒫.

#### Probability that *m* + 2 skips to *m*

The probability that a backward WF applied to a sample of size *m* + 2 results in a sample of size *m*, conditioned on a merger, is

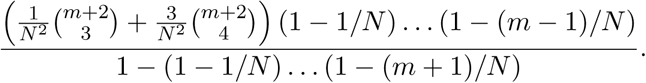

There are 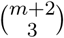 possible triple mergers and 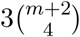 possible double binary mergers. The first factor in the numerator accounts for the probabilities of those. In both a triple merger and a double binary merger, a total of *m* parents must be chosen distinctly, which occurs with probability (1–1*/N*) *…* (1–(*m–*1)*/N*). That accounts for the second factor in the numerator. The denominator is the probability that the number of parents of *m* + 2 samples is fewer than *m* + 2 in a single backward WF step.

The above expression is simplified to obtain (9).

#### The event *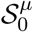*

For convenience, we restate the expression for *W*_0_ given in (10):

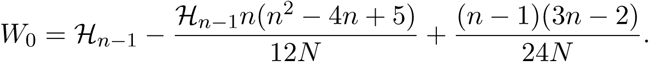

The corresponding passage in the main text defines *ℳ*_*m*_ as the event where an ancestral sample of size *m* in the genealogy of the current sample of size *n* is hit with a mutation. The expression for 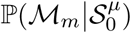

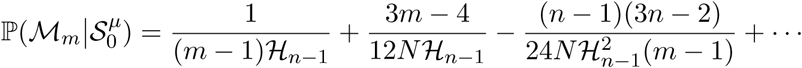

is also restated for convenience.

The probability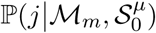, where *j* stands for *j* mutants in the current sample of *n*, is given by (22). By writing (3*m–*4) (*m*–1) as 3(*m*–1)^2^–(*m*–1), we obtain

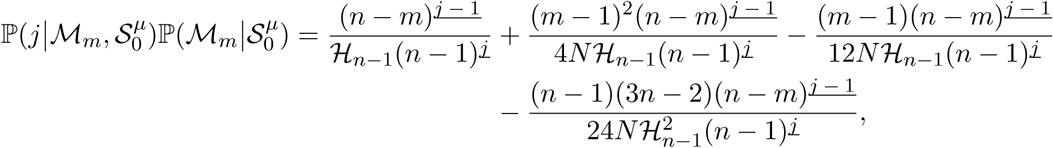

with *N* ^*-*2^ terms ignored and in the limit *µ ⟶* 0. We then have

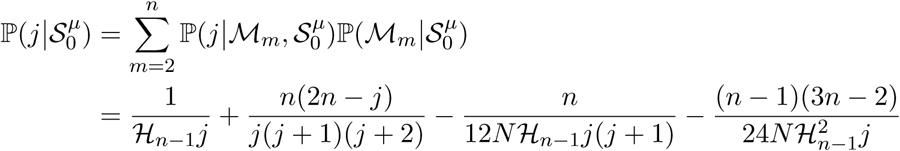

after simplification, with *N* ^*-*2^ terms ignored and in the limit *µ ⟶* 0. The simplification is effected using the following identities:

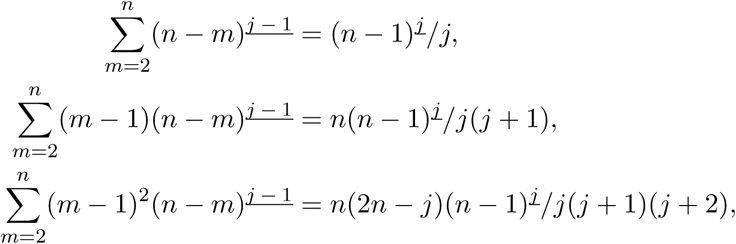

for *j* = 1, 2, *…*, each of which is easily proved by induction on *n*. Another method of proof is to begin with the difference identity (*n* + 1)^*<u>j</u>*^ *-n*^*<u>j</u>*^ = *jn*^*<u>j</u>*^.

#### The event *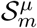*

For convenience, we recall from (11)

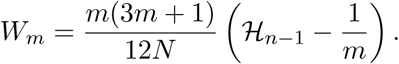

That *W*_*m*_ leads with an *N* ^*-*1^ term has the advantage that *N* ^*-*1^ terms can ignored in the rest of the 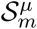 calculations. To calculate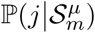, we use a shortcut that greatly simplifies the algebra. Under the condition 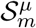 and by (8) (with the *µ*_*m*_*/*12*N* term ignored because *W*_*m*_leads with a *N* ^*-*1^ term), the probability of a mutation at *l* is proportional to 1*/*(*l -* 1) for *l ∈ {*2, *…, n} - {m* + 1*}*. Therefore the probability of a mutation at *l* under the condition 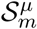 is equal to

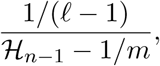

in the limit *µ ⟶*0 and with *N* ^*-*1^ terms ignored. Now for the shortcut, suppose we can ignore the WF corrections to the propagators, namely, the latter terms in the WF propagators (24a),(24b), and (24c)., We can then obtain the probability of *j* mutants in the current sample of *n* to be

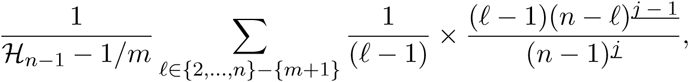

where the single mutant at *l* is propagated to *n* using the coalescent propagator (22) before summing over *l*. This expression can be simplified to get

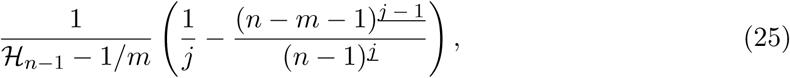

which is the probability of *j* mutants except for the corrections given by the latter terms in the WF propagators (24a), (24b), and (24c).

We will now calculate the corrections separately. First note that

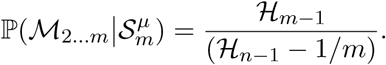

The latter terms in the WF propagators (24a), (24b), and (24c) will be activated only when the condition *ℳ*_2*…m*_ holds in addition to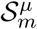.

Conditioning on *ℳ*_2*…m*_ and 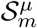, the frequency spectrum of ancestral sample of size *m* is given by

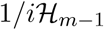

for the probability of *i* mutants, *i* = 1, *…, m–*1 (in the limit *µ →* 0 and with *N* ^*-*1^ terms neglected). To obtain the correction, this frequency spectrum must first be propagated to *m* + 2 samples using the latter terms of the WF propagators (24a), (24b), and (24c) because the condition 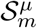 stipulates a skip form sample size *m* + 2 to sample size *m*. Propagating the probabilities to *m* + 2, we get the corrections to the probability of *i* mutants in a sample of *m* + 2 under the conditions *ℳ*_2*…m*_ and 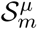 to be

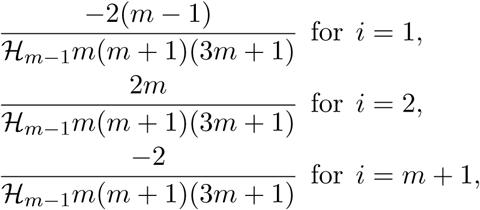

and zero for all other *i∈*{1, *…, m* + 1}–{1, 2, *m* + 1}. Multiplying these numbers with the coalescent propagator (21) with *m⟵m* + 2 and *i⟵*1, 2, *m* + 1, respectively, we get the corrections to the probability of *j* mutants in the current sample of *n* under the conditions *ℳ*_2*…m*_ and 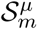 to be

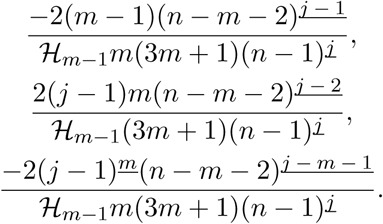

Multiplying these terms by 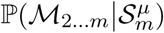 and adding to (25), we get

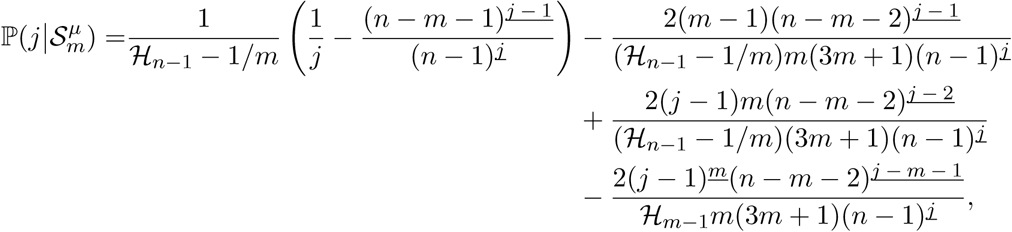

in the limit *µ ⟶*0 and with *N* ^*-*1^ terms ignored.

The sum 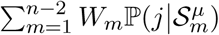 may be simplified to get

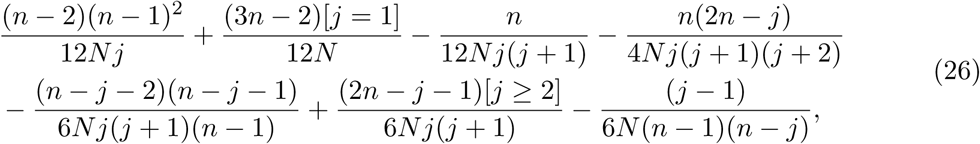

where the second line accounts for WF corrections to the coalescent propagators. The simplification uses the identities

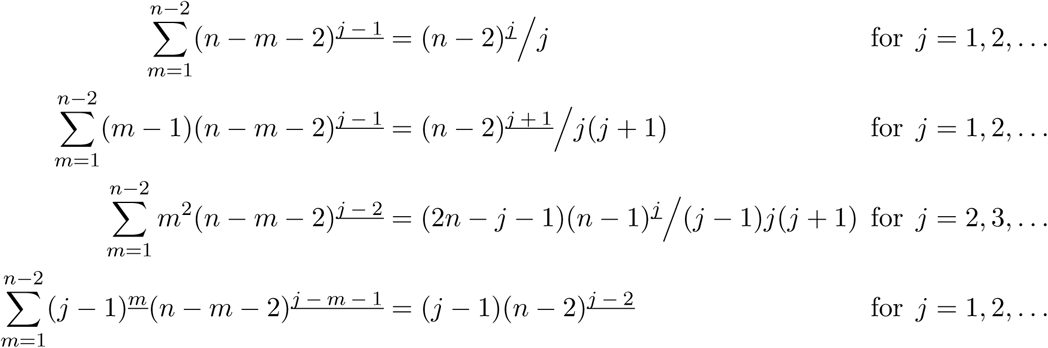

In the last identity, a^<u>b</u>^ is assumed to be 1 if b*≤* 0. All these identities may be verified by induction on *n*.

The WF sample frequency spectrum given by (2) in the limit *µ ⟶*0 and with *N* ^*-*2^ terms neglected is obtained by applying routine simplifications to

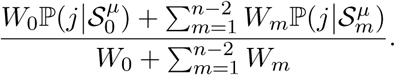

The part of the correction solely due to the difference in the way WF and the coalescent partition children between parents, which is given by (14), is obtained from the second line of (26).

If the number of samples is *n* = 3, the exact WF frequency spectrum is given by

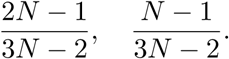

If *n* = 4, the exact WF frequency spectrum is given by

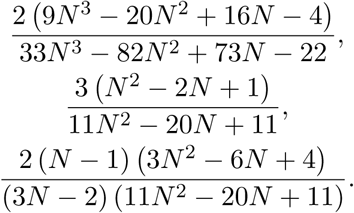

The perturbative WF frequency spectrum (2) may be checked against these exact answers.

#### Population sized samples

The coefficient of *x*^*b*^ in (exp(*αx*) *-* 1)^*a*^*/*(exp(*α*) *-* 1)^*a*^ is given by

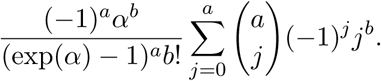

From here (17) may be derived using [Graham et al., 1994, (6.19), p. 265] to evaluate the sum.

Now we explain how to solve (15), (18), and (19) using Chebyshev polynomials. For convenience, we restate the recurrence for *g*_1_:

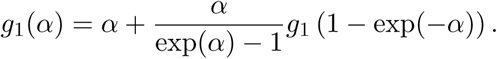

Begin with *g*_1_(*α*) = *C*_0_ + *C*_1_*α* + *C*_2_*α*^2^ + *C*_3_*α*^3^ + …and expand each term of the recurrence to obtain all terms up to the *α*^3^ term. We then obtain

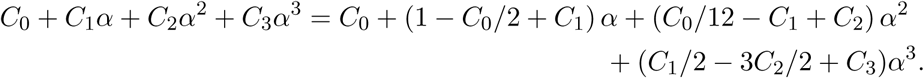

If follows that *g*_1_(*α*) = 2 + *α/*6 + *α*^2^*/*18 + …as in [Wakeley and Takahashi, 2003]. Using the same method, we get *g*_2_(*α*) = 1 –*α*^2^*/*36 + 𝒪 (*α*^3^) and *g*_*b*_(*α*) = 2*/b* + 𝒪 (*α*^3^) for *b >* 2.

To solve the recurrence for *g*_1_(*α*), we set *g*_1_(*α*) = 2 + *α/*6 + *α*^2^*/*18 + g_1_(*α*). The resulting recurrence of g_1_(*α*) is

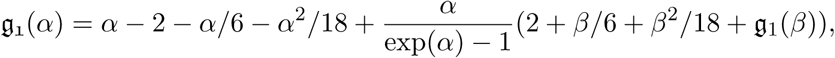

where *β* = 1–exp(–*α*). It is solved by iteration at each of 32 Chebyshev points in *α ∈* [0, 1]. The function *g*_1_(*α*) may then be obtained with 10+ digits of accuracy at any *α ∈*[0, 1] using the barycentric Lagrange interpolant [Trefethen, 2013]. The functions *g*_*b*_(*α*), with *b* = 2, 3, …, 20 are calculated using the same method.

For the functions 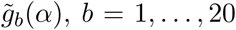, we begin with 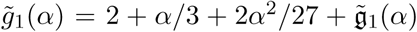,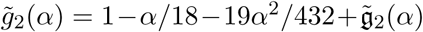 and 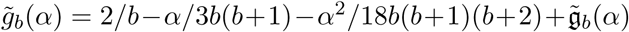 for *b >* 2. The rest of the method is the same.

To verify that 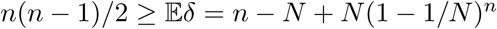,set

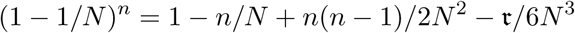

and use the Lagrange form of the Taylor series remainder to deduce τ *>* 0.

The inequality 2((1 *-* exp(*-α*))^*-*1^ *- α*^*-*1^) *>* 1 is equivalent to

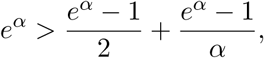

which is proved by verifying that the series for the left hand side majorizes the series for the right hand side.

To show that 2*N* (*n-m*)*/*(*n-*1) *< Nα* when *n* = *Nα* and *m* = *N* (1 *-*exp(*-α*)), first observe the inequality follows from 2(1*-m/n*) *< α* for *N* large. Now 2(1*-m/n*) *< α* is equivalent to *e*^*-α*^*<* 1 *-α* + *α*^2^*/*2, which can be verified using the Lagrange form of the Taylor series remainder.

### Discussion

Suppose the sample size is *αN* with the parental population size being *N* as usual. Conditional on an individual of the parental generation being a parent of one of the samples and assuming *N* large, its number of children (among the samples) is given by the generating function (exp(*αx*)*-*1)*/*(exp(*α*)*-*1). It follows that the expectation of the number of children is *α* and the variance is

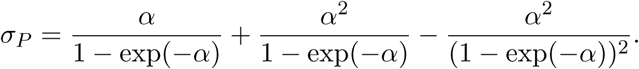

If the *αN* children are split among their parents according to the Kingman partition distribution, the generating function for the number of children is *γx/*(1 *-* (1 *- γ*)*x*) with *γ* = (1 *-* exp(*-α*))*/α*. The expectation is again *α* and the variance is

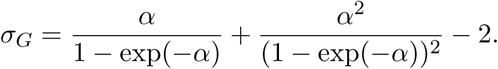

One may verify that *σ*_*G*_*> σ*_*P*_ by plotting a graph. Alternatively,

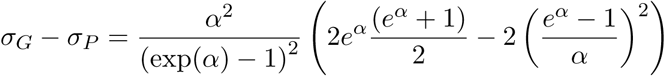

must be positive because the power series of both *e*^*α*^and (*e*^*α*^ + 1)*/*2 majorize the power series of (*e*^*α*^ − 1)*/α*.

Intuitively, we expect *σ*_*G*_*> σ*_*P*_ because the geometric distribution has exponential decay, whereas the Poisson distribution has super-exponential decay.

## Footnotes

1 Wright’s result is the same as the modern coalescent estimate of the expected size of the genealogy, with 1.8 being his approximation to *e*^γ^, where *γ* is Euler’s constant. Wright’s 2*N* is the same as our *N* and his *μ* is the same as our 2*μ*. We have modified his formulas accordingly.

